# Bats of a feather: Range characteristics and wing morphology predict phylogeographic breaks in volant vertebrates

**DOI:** 10.1101/2024.02.11.579809

**Authors:** Sydney K. Decker, Kaiya L. Provost, Bryan C. Carstens

## Abstract

Intraspecific genetic variation and phylogeographic structure can be influenced by factors such as landscape features, environmental gradients, historical biogeography, and organismal traits such as dispersal ability. Since deep genetic structure is often considered a precursor to speciation, identifying the factors that are associated with genetic structure can contribute to a greater understanding about diversification. Here, we use repurposed data to perform a global analysis of volant vertebrates (i.e., bats and birds) to estimate where intraspecific phylogeographic breaks occur and identify the factors that are important predictors of these breaks. We estimate phylogeographic breaks using Monmonier’s maximum difference barrier algorithm and conduct a Random Forests analysis using the presence of a phylogeographic break as a response variable. In bats, phylogeographic breaks are concentrated in biodiversity hotspots while breaks estimated in bird species are more widespread across temperate and tropical zones. However, for both clades geographical features such as maximum latitude, measures of wing morphology, and organismal traits associated with feeding ecology were found to be important predictors of phylogeographic breaks. Our analysis identifies geographical areas as wells as suites of organismal traits that could serve as a starting point for more detailed studies of biodiversity processes.

## Introduction

Intraspecific genetic variation is a featured aspect of various models of speciation (Lande 1980; Orr 1996; Nosil 2012). One central tenet of evolutionary theory is that natural selection acts on variation within species to drive diversification (e.g., Grant 1981). For this reason, entire disciplines have been devoted to characterizing intraspecific genetic variation and identifying the landscape features that may influence it (e.g., Avise 2000; Manel and Holderegger 2013). While synthesis has been impeded by confounding factors such as life history traits, environmental variation, and biogeographic history, such work has resulted in the accumulation of millions of sequences on databases such as NCBI GenBank (e.g., Garrick et al., 2015). Coupled with other data, such as specimen collection records from natural history collections (e.g., iDigBio), data aggregation portals (e.g., GBIF), and collections of phenotypic data (e.g., MorphoBank), understanding the determinants of intraspecific genetic diversity broadly across taxonomic groups now appears to be an attainable goal (Barrow et al. 2021).

One key phenomenon that is likely to influence the development of intraspecific genetic variation is dispersal, particularly intrinsic and extrinsic factors that limit the ability of individuals of a species to move throughout the landscape. The lack of dispersal can lead to the accumulation of genetic differentiation due to genetic drift and other forces (Slatkin 1987), and dispersal limitation has been identified as a key factor in biological processes ranging from speciation rates (e.g., Polato et al. 2015) to community assembly (e.g., Pigot and Tobias 2015). Research into empirical systems has indicated that landscape features such as mountains (e.g., Kane et al. 2019), rivers (e.g., Myers et al. 2020), and environmental gradients (e.g., Sabastián et al. 2021) can limit dispersal and lead to the formation of intraspecific genetic variation.

Long-term limitation of gene flow between populations, with or without a physical barrier to gene flow (e.g., Irwin 2002), can result in phylogeographic structure. Phylogeographic structure is often described using summary statistics describing genetic variation in populations (e.g., F_ST_), identifying population genetic clusters (e.g., PCA), or by testing for spatial autocorrelation.

Considerably less effort has been devoted to theory and methodology for estimating *where* barriers to gene flow occur. One notable exception is Monmonier’s algorithm (Monmonier 1973), which can be applied to infer spatial genetic structure by converting the genetic data into a network, identifying the edges of the network that are associated with the highest degree of genetic differentiation, and projecting the maximum difference boundary onto a spatial connection network (e.g., Manni et al. 2004). The linear boundary produced by Monmonier’s algorithm is placed on a map without any input from external information such as geospatial or environmental layers, making the algorithm well-suited for macrogenetic investigations which seek to identify the factors that influence genetic structure. Since geographic distance is typically correlated with environmental distance and landscape features (e.g., Wang 2013, Pelletier and Carstens 2018), any inference method that incorporates geographic distance among samples will likely be influenced by these same factors. Monmonier’s algorithm circumvents this problem and enables environmental distance and landscape features to be evaluated as potential factors which may contribute to the formation of phylogeographic breaks.

Due to the perception that volant species have the capacity for long distance dispersal, phylogeographic breaks may be more likely to correspond to environmental conditions, differences in food resources, and overall habitat suitability than to physical and geological barriers. In bats, patterns of phylogeographic structure vary across species on a global scale but are thought to be influenced by factors including social structure, mating behavior (e.g., autumnal swarming), migration patterns, habitat connectivity, ecological gradients, and geographic barriers (Moussy et al. 2013). In a recent review, Hernández-Canchola et al. (2021) noted that Quaternary climate fluctuations, geographic features such as oceans and barriers, and ecological processes such as niche differentiation were the most referenced contributors to phylogeographic structure in bat species. In birds, feeding ecology and annual dispersal were the strongest predictors of high diversification rates (Phillimore et al. 2006), suggesting that species ecology plays a large role in diversification and speciation. Morphological proxies of dispersal show a nuanced response with respect to diversification (Claramunt et al. 2012, Tobias et al. 2020). Social behavior, particularly song, is also thought to be important at maintaining diversity across the landscape (Uy et al. 2018)

To assess global phylogeographic patterns in volant vertebrates and the factors that are important in predicting them, we repurposed georeferenced mitochondrial sequence data available from *phylogatR*, a database that aggregates geographic information from specimen repositories such as GBIF and sequence data from GenBank and BOLD in a common framework (Pelletier et al. 2022). While challenges related to data acquisition, quality control, and unevenness can be substantial and should be accounted for (Leigh et al. 2021; Pelletier et al. 2022), we apply an automated analysis pipeline modified from previous investigations (Pelletier & Carstens 2018; Barrow et al. 2021; Parsons et al. 2022) to estimate phylogeographic breaks in hundreds of bird and bat species. Bats and birds were chosen for the study as they are both volant vertebrates with high ecological diversity and are generally thought to be good dispersers. Additionally, birds tend to be well studied in comparison to bats, and as such we hoped that the additional Avian data would corroborate models of intraspecific genetic structure in bats. We then use machine learning to identify traits that are predictive of phylogeographic breaks in these taxa.

## Methods

### Data Processing

Data for global Chiroptera and Aves species were downloaded *phylogatR* (phylogatr.org; Pelletier et al. 2022). Custom R scripts (available on https://github.com/skdecker/PhylogeographicBreaks) were developed to further filter occurrence and sequence data with the goal of removing data that were likely to be inaccurate. To accomplish this, we removed geographic data that fell outside species range maps. Coordinates were cleaned based on IUCN range maps for bats (IUCN 2022) and BirdLife range maps for birds (BirdLife International 2021) with a 1° (approximately 110 km) buffer to account for animal movement and range map inaccuracies. To detect poor quality genetic data, sequence data were processed to remove sequences with greater than 20% missing sites from alignments, as such data were likely to lead to failures in downstream analyses. We also removed sequences that exhibited high genetic distance from other sequences in the alignment (>5% in birds and >10% in bats) because data such as these might be indicative of hidden species (e.g., Parsons et al. 2022) or misidentified individuals. These thresholds are based on the distribution of average intraspecific genetic distances in our data (supplementary figure S1) and historical quantifications of mitochondrial genetic distances typical of biological species (Baker and Bradley 2006). Due to the historical use of mtDNA in phylogeographic studies and for ease of comparison across species, we relied exclusively on mtDNA for these analyses. Since over 86% of the sequence alignments downloaded from *phylogatR* for birds and bats were from mitochondrial genes, this also allowed us to include the largest possible sample size. Species with 15 or more sampled individuals in the alignment and at least 3 unique localities after processing were included in downstream analyses. In species with data for multiple genes, the gene with the highest number of sequences was used.

### Estimating Phylogeographic Breaks

In order to identify species traits that are associated with the formation of genetic structure, we first need to identify species that exhibit spatial population genetic structure. A quantitative approach was preferred since this could be applied across hundreds of species and would not be biased by the interpretations of individual researchers, as would be the case if we conducted a synthetic literature review. We found it expedient to apply a method that does not require *a priori* division of samples, such as calculating F_ST_ values. We did not use a test of genetic isolation by distance (Wright 1943) due to the anticipated correlation between geographic and environmental distance (e.g., Dillon 1984; Lee and Mitchell-Olds 2011) and the association of genetic isolation by distance with other factors such as latitude (Pelletier & Carstens 2018). Rather, Monmonier’s algorithm (Monmonier 1973) was applied to estimate phylogeographic breaks.

Monmonier’s algorithm was implemented through the optimize.monmonier function in the R package *adegenet* v2.1.4 (Jombart 2008; Jombart and Ahmed 2011). Monmonier’s algorithm is incompatible with duplicate localities, therefore a small amount of noise was added to duplicate coordinates with the R function jitter (base R, v4.1.1; R Core Team 2021). The input data for the function is a connection network created from jittered sample coordinates and a matrix of genetic distances. For our analyses, we used Euclidean distances for the genetic data and geographic connection networks built using Delaunay triangulation using the chooseCN function in *adegenet*. Unlike the standard monmonier function, optimize.monomonier tries several starting points within the connection network and returns the best boundary, avoiding single strong local differences (e.g., Dupanloup et al. 2002). The number of tries used in the algorithm was proportional to the number of individuals for each species. A maximum of 10 tries was used for species with greater than 100 sequences for computational efficiency. The results of the function are output as a list of coordinates comprising discovered boundaries. These returned coordinates were plotted as lines on a map with the sampling localities and the jittered sampling coordinates for each species individually to manually check for phylogeographic breaks.

Species were assigned to one of two categories (i.e., ‘no phylogeographic break’ or ‘phylogeographic break’) based on whether the algorithm returned a break that clearly divided the sampling range, isolating individuals/populations from one another. In cases where the presence of a phylogeographic break was ambiguous, the local distances plot from the Monmonier’s algorithm run was evaluated: a sharp decrease in the plot suggests a break is present while a steady cline suggests that no break is present. This assignment was used as a binary response variable in the following random forests classifier.

### Traits

Once genetic data were acquired and phylogeographic breaks were estimated, an organismal trait dataset (supplementary data sheets 1,2) was compiled from published, open-source datasets. For bat traits, we used PanTHERIA (Jones et al. 2009), COMBINE (Soria et al. 2021), and Crane et al. 2022. Some missing values for wing morphology metrics were supplemented by a literature search (supplementary data sheet 1). AVONET (Tobias et al. 2022) was used for bird traits, following the BirdLife taxonomic system. The datasets consisted of geographic (e.g., maximum latitude), taxonomic (family/order), morphological (e.g., mass), ecological (e.g., trophic level), and environmental (e.g., habitat) traits for each species. Because Random Forests cannot be trained with missing data, traits with more than 40% missing data across species or species with more than 40% missing trait values were removed from the analysis and missing values for remaining traits were imputed using the R package *mice* v3.15.0 (van Buuren and Groothuis-Oudshoorn 2011). For bats, the taxonomic Family was used as a predictor in the imputation.

Penone et al. (2014) found little bias in methods such as *mice* for imputing missing trait variables, especially when taxonomic information is used as a predictor. Imputation was conducted over 500 iterations. For each variable with imputed data, the imputed values were checked for bias against observed data with the densityplot function (supplementary figure S2). Missing data were only imputed for the bat trait dataset as AVONET reports inferred values from the closest relative in cases of missing values (Tobias et al. 2022).

### Random Forest Classifier

To identify traits important in predicting whether a species will exhibit a phylogeographic break, we built a Random Forests classifier with the R package *caret* v 6.0-88 (Kuhn 2021) to identify which of the trait data were responsible for a particular species being classified in either category. To improve fit of the RF model, we used recursive feature elimination (RFE) to test subsets of between 1 and 25 predictor variables with 25-fold cross validation and 5 repetitions with the rfe function from *caret*. Additionally, correlation of predictor variables was assessed with cor function from the R package *stats* v4.1.1 (R Core Team 2021) and the package *corrplot* v0.92 (Wei and Simko 2021; supplementary figures S3, S4). Predictor variables identified by recursive feature elimination were retained for the classifier. We split data into training (85%) and testing (15%) datasets, and 2000 decision trees were used in training the classifier with 5-fold cross validation. Out of bag (OOB) error rate, within-class error, and variable importance metrics were averaged over 50 random seeds of RF and accuracy of the final models were assessed using the testing datasets.

## Results

### Results of Data Processing

Genetic and locality data for 383 species of bats and 1971 species of birds were downloaded from *phylogatR*. After filtering for missing data, high sequence distances, occurrences outside of published geographic ranges, and fewer than 15 sequences per alignment, we retained 126 bat species and 214 bird species. Though there were substantially more bird species than bat species in the original dataset, a larger proportion of bird species (∼82%) were excluded prior to filtering steps due to low number of sequences per species. The average length of the alignments was 688 base pairs (bp) for bats and 787 bp for birds.

### Results of Monmonier’s Algorithm

A conspicuous phylogeographic break was estimated in 68 bat species (54%) and 95 bird species (44.4%) using Monmonier’s algorithm. In bats, these phylogeographic breaks were primarily clustered in areas of high diversity and sampling, specifically southern Central America and northern South America and southeast Asia (figure 1). Breaks estimated in birds were more widespread across temperate zones, South America, and southeast Asia (figure 1). Longer, transcontinental and intercontinental phylogeographic breaks estimated for birds are likely explained by the larger geographic ranges of birds in our dataset (mean: 11,803,801 km^2^) compared to range size of bats included (6,930,099 km^2^).

**Figure 1:**
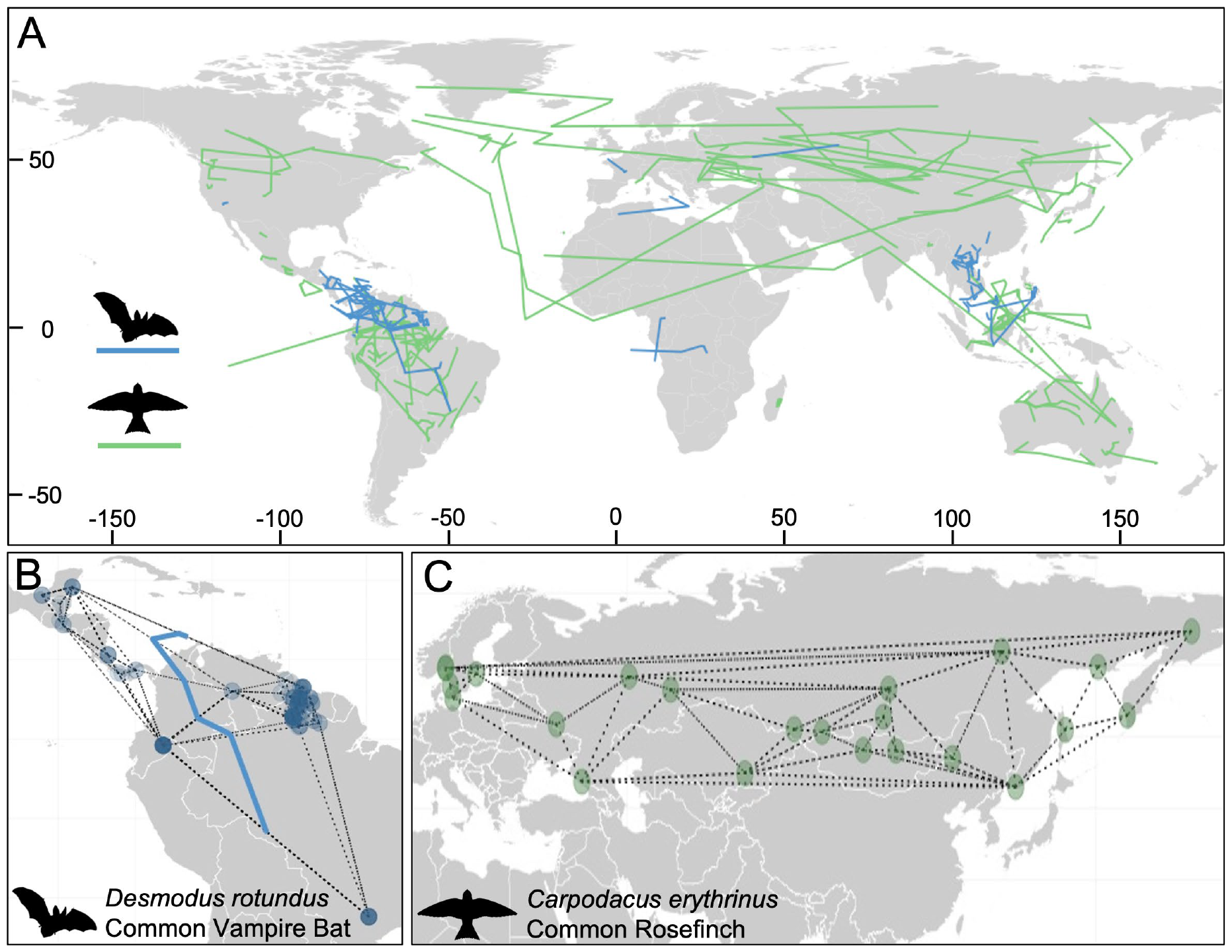
A) Map of phylogeographic breaks estimated by Monmonier’s algorithm for global bats (blue) and birds (green); B) Example of species in which a break (thick blue line) was estimated, *Desmodus rotundus*, with connection network between occurrences C) Example of species determined to not contain a phylogeographic break, *Carpodacus erythrinus*.

### Results of Random Forest Classifier

Missing data were imputed for the bat trait dataset and density plots indicated that bias in imputed values was minimal. Recursive feature elimination was used to decrease the number of predictor variables and simplify the models used. In the bat dataset, RFE indicated that 15 predictor variables should be included in the classifier after removal of highly correlated non-morphological variables (supplementary data sheets 3,5). For birds, the best model included 23 variables. We assessed preliminary versions of the trained models and found variation in accuracy and in which variables were most important in predicting if species exhibited a phylogeographic break, so 50 independent classifiers were trained for each dataset.

Across 50 random seeds, the out of bag (OOB) error rate for the bat dataset classifier ranged between 0.3001–0.3974, with an average accuracy of 66.00% (table 1, supplementary data sheet 12). The classifier for the bird dataset had an average accuracy of 62.48%, with OOB error rate ranging from 0.3397–0.4089 (table 1, supplementary data sheet 13). The bat classifier was more accurate in predicting bat species that contained a phylogeographic break (class error 0.2313) compared to bat species that did not exhibit a break, while the bird classifier more accurately predicted species that did not exhibit a phylogeographic break (class error 0.2965) than those that did (table 1). The bat classifier had higher sensitivity and precision, indicating that the model slightly over-predicted that species would contain breaks and the bird model had higher specificity, indicating a tendency to predict that species would not exhibit phylogeographic breaks (table 1). This result is expected considering the slightly uneven distributions of the response variable in the bat and bird datasets (Adam et al. 2014).

**Table 1.** Statistics for the trained Random Forests models.

|  | OOB | Class (N) | Class (Y) | Sensitivity | Specificity | Precision |
| --- | --- | --- | --- | --- | --- | --- |
| Bats | 0.3400 | 0.4661 | 0.2313 | 0.7540 | 0.5400 | 0.6798 |
| Birds | 0.3752 | 0.2965 | 0.4745 | 0.5657 | 0.7047 | 0.6245 |

### Important Predictors

In bats, the ten top predictor variables as measured by Mean Decrease in Accuracy (MDA) included occurrence area, cryptic diversity predicted, latitude length, maximum latitude, mountains within range, brain mass, minimum longitude, wingspan, wing loading, and carnivory (figure 2a). Occurrence area and latitude length were significantly larger in species estimated to contain a phylogeographic break compared to those without a phylogeographic break (p=0.002, p=0.0002, respectively). Species with a phylogeographic break also exhibited significantly higher maximum latitudes (p=0.006), smaller brains (p=0.048), and shorter wings (p=0.045).

**Figure 2.**
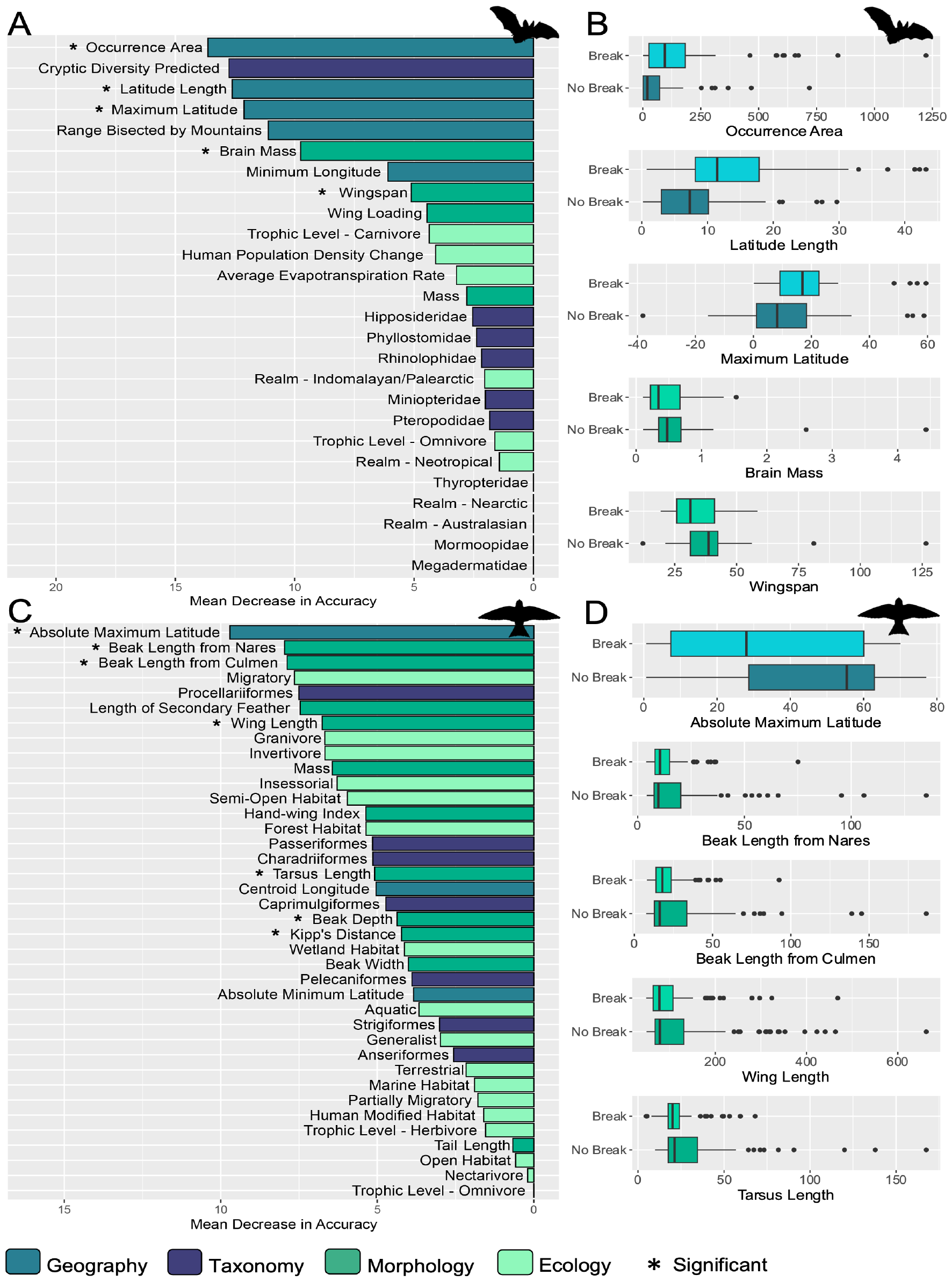
(a) Predictor variable importance by mean decrease in accuracy (MDA) for bats, averaged across 50 independently trained classifiers; (b) boxplots for significant variables in the bat dataset; (c) predictor variable importance by MDA for 50 independently trained bird model; and (d) boxplots for five significant variables in the bird dataset.

Wing loading was also lower, on average, in bat species estimated to contain a phylogeographic break, though this relation was not statistically significant (p=0.2001; supplementary table S1).

In birds, absolute maximum latitude, beak length from the nares, total beak length along the culmen, status as migratory, order Procellariiformes, length of the first secondary feather (a proxy for wing width; figure 3b), wing length, granivore and invertivore trophic niches, and mass were the ten most important predictors (figure 2c). Bird species with a phylogeographic break exhibited absolute maximum latitude closer to the equator (p=0.0002) and a smaller minimum latitude (p=0.021). Species with breaks also had shorter beaks both in length from the nares (p=0.022) and in total length along the culmen (p=0.014) and shorter wings (p=0.028).

**Figure 3.**
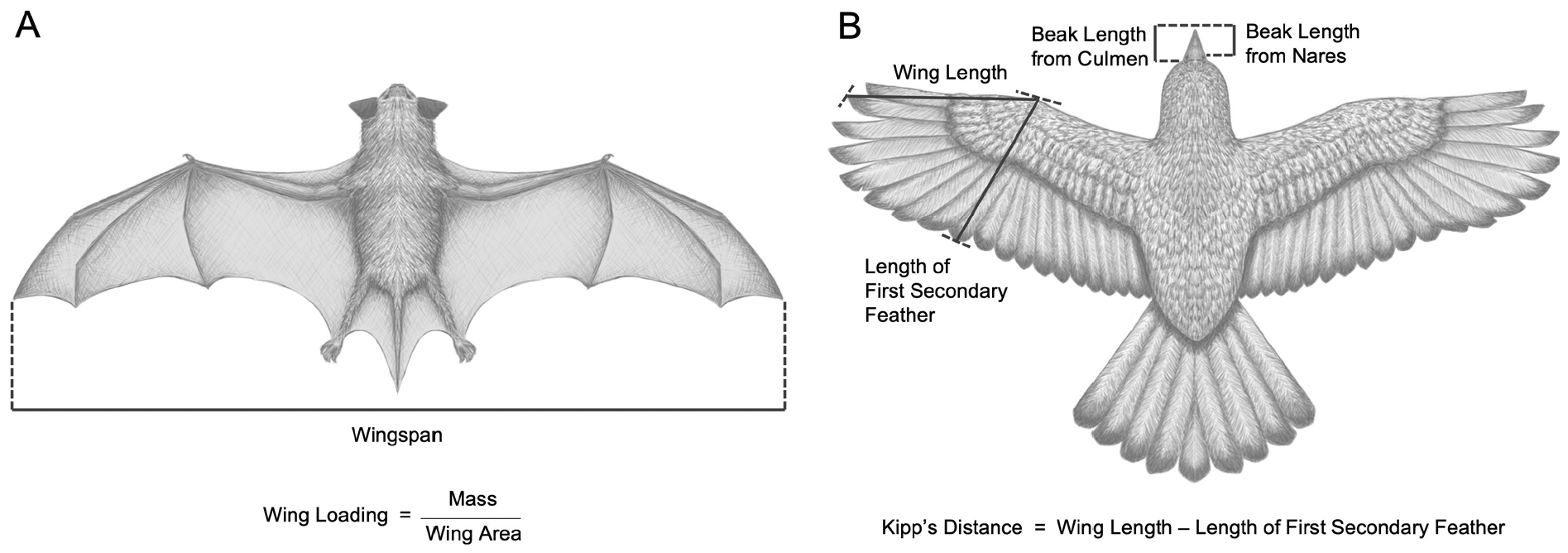
(a) Bat illustration with wingspan measurement indicated and equation for wing loading; (b) Bird with important morphological measurements indicated and the equation for Kipp’s distance.

Though not found in the top ten most predictive variables, species estimated to contain a phylogeographic break had shorter tarsus length (p=0.002), more shallow beak depth (p=0.016), and shorter Kipp’s distance (p=0.02732). Though not statistically significant, a trait considered to be important to dispersal ability in birds (hand-wing index; Claramunt et al. 2012) was on average lower in bird species with a phylogeographic break (p=0.167; supplementary table S2).

## Discussion

Consistent with previous investigations into population genetic structure (e.g., Martin and McKay 2003; Pelletier and Carstens 2018), many top predictors of phylogeographic breaks in bats and birds include range characteristics such as occurrence area and measures of latitude. In our study bird species that contained phylogeographic breaks exhibit maximum and minimum latitudes closer to the equator, consistent with the view that tropical bird species are expected to be more sedentary, have decreased dispersal ability (Sheard et al. 2020), and have more genetic population structure (e.g., Brawn et al. 1996). Bat species with a higher maximum latitude more often exhibit phylogeographic breaks, a pattern likely driven by low sampling in temperate zones and the concentration of breaks found above the equator in southern Central America and southeast Asia (supplementary figure S5). Additionally, the presence of physical barriers such as a mountain range can still play an important role in structuring genetic diversity even in volant vertebrates as such barriers were a top predictor of phylogeographic breaks in bats. But landscape is not destiny, as some species of both birds and bats seem to have found a way to preserve genetic connectivity across their range. The organismal traits that facilitate this connectivity are likely related to those required by powered flight, notable features of the wing that influence flight performance and efficiency.

Most of the organismal traits that were found to be important predictors of phylogeographic breaks in bats and birds are associated with wing morphology (Figure 3). Wing loading (i.e., the surface area of a wing relative to the mass of the flying object) and wing aspect ratio (i.e., the ratio of the wing span to the wing width) are likely to be important because these factors influence the efficiency of flight in flying objects that range in size from *Drosophila* to the Airbus A380. In bats, higher wing loading is positively correlated with flight speed and distances (Norberg and Rayner 1987) whereas high aspect ratios are found in species with long, thin wings that exhibit lower drag and increased aerodynamic efficiency (Norberg 1995). Both of these characteristics are likely correlated with higher dispersal ability, larger geographic ranges (e.g., Luo et al. 2019), and greater potential to traverse physical barriers in the landscape (Burns and Broders 2014). Therefore, bat species that exhibit high wing loading and longer wing morphologies are more likely to exhibit genetic connectivity across their ranges and less intraspecific genetic structuring (Burns and Broders 2014, Taylor et al. 2012). Similarly, in birds increased dispersal ability with more efficient long-distance flight is associated with elongated wings with higher hand-wing index, a measure similar to aspect ratio (Claramunt et al. 2012).

Our results are consistent with findings that bats and birds with wing morphologies optimized for more efficient flight exhibit fewer phylogeographic breaks, possibly due to increased dispersal ability and genetic connectivity across their geographic ranges.

Wing shape is also implicated in bat and bird feeding ecology as determinants of how species can navigate their environment to forage (e.g., Norberg and Rayner 1987, Sheard et al. 2020). In birds, habitat specialization and diet specialization are interrelated (Rief et al. 2015). We suspect that this is also true of bats, although we cannot identify a comparable study. However, it is well established that habitat variation is associated with dietary differences within species (e.g., Clare et al. 2011), that habitat influences the guild structure of bat communities (e.g., Rocha et al. 2018), and that wing shape influence both the extent of and types of specialization of feeding ecology (e.g., Fenton 1982). In both taxonomic groups, trophic categories were included in top predictors of phylogeographic breaks and in birds, two measures of beak length were important predictors. Avian cranial morphology is usually associated with diet in birds, especially the beak (Soons et al. 2015), though this may be constrained genetically (Bright et al. 2016; Tokita et al. 2017). Bird beaks are important for other functions including thermoregulation (Greenberg et al. 2012), communication (Huber and Podos 2006), and nest building (Sheard et al. 2023). Wing morphology in passerines is also associated with diet, particularly as it relates to foraging behavior and food availability (Sheard et al. 2020). Variation in genetic differentiation across biogeographic barriers in South American bird species is also explained by foraging stratum (Bruney and Brumfield 2009), suggesting a considerable interaction between feeding ecology and genetic diversity.

Phylogeographic breaks are not necessarily indicative of species-level diversity, as phylogeographic patterns can be caused by social structure or sex biased dispersal (e.g., Dávalos and Russell 2014), historically allopatric populations (e.g., Fleming et al. 2010), limited dispersal, or small population sizes (Irwin 2002). Some phylogeographic breaks estimated here correspond to boundaries of named subspecies, for example the eastern and western subspecies of the North American Wilson’s Warbler (*Cardellina pusilla*; species specific results available on https://github.com/skdecker/PhylogeographicBreaks). Nevertheless, phylogeographic structure and intraspecific genetic diversity is positively associated with speciation rates in some taxa, for example in New World birds (Harvey et al. 2017). Furthermore, a large portion of phylogeographic breaks estimated here occur in the tropics, indicating that within species genetic differentiation follows similar latitudinal gradients as intraspecific genetic diversity (e.g., Smith et al. 2017; Fonseca et al. 2023), predictions of cryptic species (Parsons et al. 2022), and species richness (e.g., Gaston 2000, Mittelbach et al. 2007).

While repurposed, single locus, mitochondrial data are often not representative of overall patterns of population genetic structure (Bazin et al. 2006, Galtier et al. 2009), extensive data quality control steps and established automated analysis pipelines allows initial investigations of global biodiversity patterns (e.g., Barrow et al. 2020). The use of Monmonier’s algorithm to identify phylogeographic breaks has been fairly limited despite it likely being a legitimate estimate of population structure. For example, Dupanloup et al. (2002) compared Monmonier’s algorithm with spatial analysis of molecular variance (SAMOVA) and found that the algorithm performed better than SAMOVA in finding genetic breaks of simulated single locus data.

However, across different methods of phylogeographic break estimation, isolation by distance (IBD) can be misidentified as a break if sampling is sparse across the range of a species (Templeton et al. 1995; Irwin 2002).

Once phylogeographic breaks were estimated, 50 independent RF classifiers were trained due to low accuracy and instability of variable importance across preliminary models, likely caused by the use of small datasets. It was our intention that analysis of phylogeographic breaks in birds would yield a more accurate model due to increased availability of data compared to bats, however over 89% of bird species in the original download from *phylogatR* were removed by our data quality control filters. To verify that low model accuracy was due to small sample size and intrinsic variation, we trained a second RF classifier using IBD, a more widely used metric of how intraspecific genetic diversity is structured across the landscape, as the response variable against the trait datasets (supplementary material). The models trained with IBD also exhibited low accuracy (66.67% for the bat classifier, 60.88% for birds; supplementary table S3), this secondary analysis suggests that low accuracy in preliminary models is not due to the phylogeographic break response being uninformative. Though overall accuracy of both bat and bird models was still relatively low, averaging across multiple independent classifiers allowed us to have more confidence in the measures of predictor variable importance. Because RF models predict a complex of traits that interact to affect the response variable, measures such as MDA can help tease apart these relationships to infer biological meaning of influential traits.

Our investigation joins a growing body of research, described as “macrogenetics” (Blanchett et al. 2017) or “automated comparative phylogeography” (Gratton et al. 2017), that relies on the collection and reanalysis of genetic data that were originally collected for investigation into a single species. Facilitated by the accession of these data in databases such as NCBI GenBank (Benson et al. 1993) and the development of data aggregators such as phylogatR (Pelletier et al. 2022), researchers now have the capacity to ask in-depth questions about genetic diversity in a particular clade (e.g., Manel et al. 2020; Theodoridus et al. 2021; French et al. 2023) or to investigate patterns across clades on a global scale (Miraldo et al. 2016; Fonseca et al. 2023).

Macrogenetic investigations are subject to important questions about scale and overinterpretation (Paz-Vilas et al. 2021; but see Millette et al. 2021) and it is important to recognize the limitations of single locus data (Knowles 2009; Edwards and Bensch 2009). Given these difficulties and limitations, we recognize that macrogenetic investigations enable novel synthesis of biodiversity data. Our data were drawn from databases containing genetic, environmental, geographic, and organismal trait data. These data can be analyzed synthetically using machine learning or other AI approaches (e.g., Carstens et al. 2018; Parsons et al. 2022; Yang et al. 2023). Our study, conducted using data from two clades of volant vertebrate animals, indicates that feeding ecology is likely a much more important influence on intraspecific genetic structure than previously appreciated. It could be that this result is due to the unique dispersal capacity of the focal clades, or it may be that this holds true for non-volant vertebrates. In either case the question can be asked.

## Conclusions

Volant vertebrates are not equal in their dispersal capabilities. Our macrogenetic investigation and random forest analysis suggested that species which are capable of flying long distances and which have long wings that are optimized for efficiency at the expense of maneuverability are less likely to contain intraspecific genetic structure. Phylogeographic breaks tend to be found in species with shorter wings; these are optimized for maneuverable flight at the expense of efficiency over longer distances. Since species rich clades in bats (i.e., Vespertilionidae) and birds (i.e., Passerines) generally contain species do not have these wing characteristics, which suggests that wing shape may play a role in diversification rate shifts within Order Chiroptera and Class Aves. While diversification rates appear to be influenced by wing morphology in moths (Aiello et al. 2021) and there has been some exploration of these factors within Class Aves (e.g., Kennedy et al. 2016), a comprehensive phylogenetic investigation of diversification rates and wing morphology in Chiroptera and Aves is needed.

## Acknowledgements

We thank members of the Carstens lab for their helpful comments on this work. Support for this work was provided by the National Science Foundation (NSF) (DBI-1661029 and DBI-1910623 to B.C.C.) and the Ohio Supercomputing Center (PAA1174).

## Data Availability Statement

Code and source data, including DNA sequence alignments, trait data, and analysis files, used in the manuscript are available at https://github.com/skdecker/PhylogeographicBreaks. All other data are provided in supplementary material.

