## Supplemental Files for "Bats of a feather: Range characteristics and wing morphology predict phylogeographic breaks in volant vertebrates"

**Supplementary Material**

### Supplementary Methods and Results

Isolation by Distance Analysis

In addition to building a predictive model using phylogeographic breaks as a response variable, we also tested a different summary of phylogeographic structure, isolation by distance (IBD), against our predictor traits. IBD was calculated for each species using mantel tests with the function mantel.randtest in the R package *ade4* v1.7-19 (Dray and Dufour 2007) with 500 replications. Results of the IBD analysis were coded as significant IBD ($p<0.05$) or nonsignificant IBD ($p>0.05$) in a binary RF classifier analysis. Data were split into training (85%) and testing (15%) datasets, and 5000 decision trees were used in training the data with 5-fold cross validation. Out of bag (OOB) error rate, within-class error, and variable importance metrics were computed for the trained model and accuracy of the model was assessed using the testing dataset.

IBD Results

Signatures of significant IBD were found in 89 of bat species in the study (70.6%) and 128 bird species (59.8%). The trained RF classifier for the Chiroptera data showed improved overall accuracy (71.4%) compared to the classifier predicting a phylogeographic break (66%), however there is high class imbalance and low precision (33.3%) indicating a tendency to almost always predict the major class. For the Aves data, the IBD classifier was slightly less accurate overall (59.4%) compared to the phylogeographic break classifier (62.48) with low specificity (25%). Though not identical to predictors found to be important in predicting phylogeographic breaks, similar traits were found to be important in predicting isolation by distance (Figure S6). Range characteristics, traits relating to feeding ecology, and morphological traits associated with dispersal ability were top predictors of IBD in birds and bats (Figure S6).

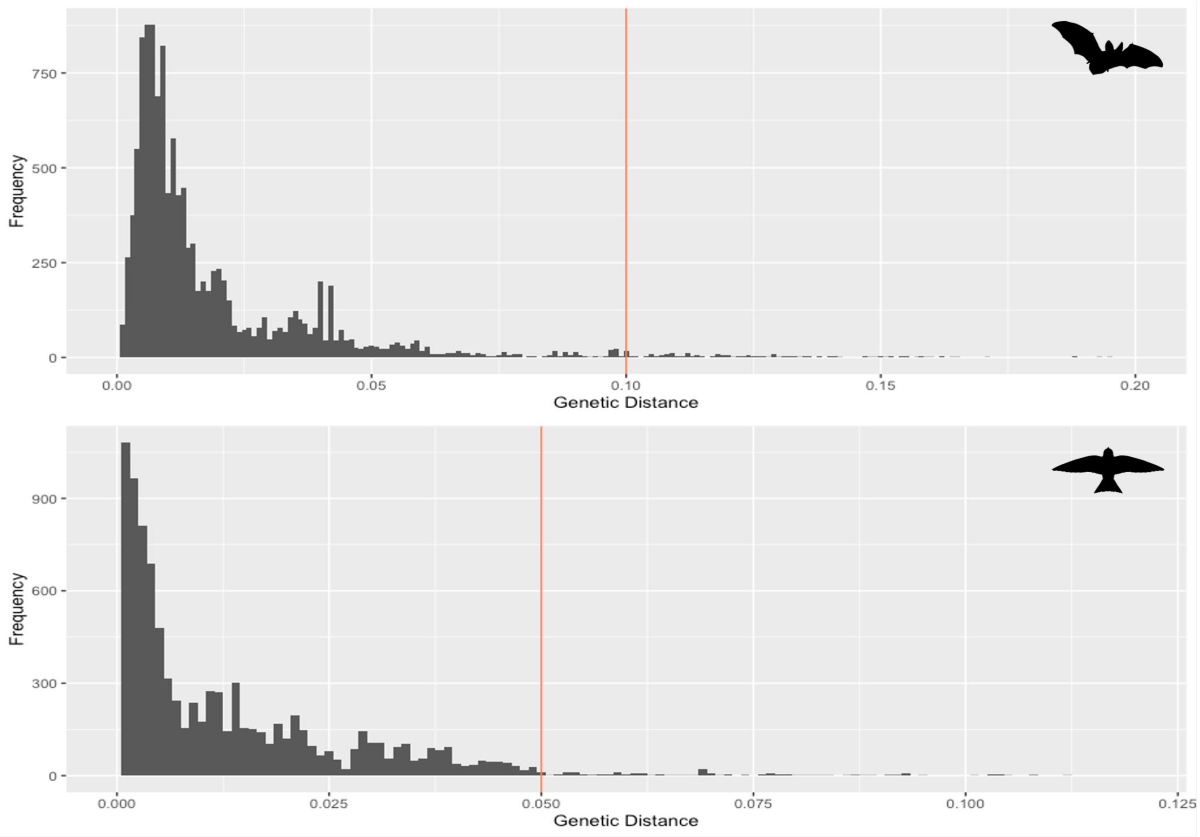


**Figure S1.** Frequency histogram of genetic distances of sequences in *phylogatR* dataset for bats (top) and birds (bottom) used to help determine cutoff for genetic distance to decrease inclusion of misidentified individuals. Chosen values for distance limit indicated by colored vertical line: 0.10 for bats, 0.05 for birds.

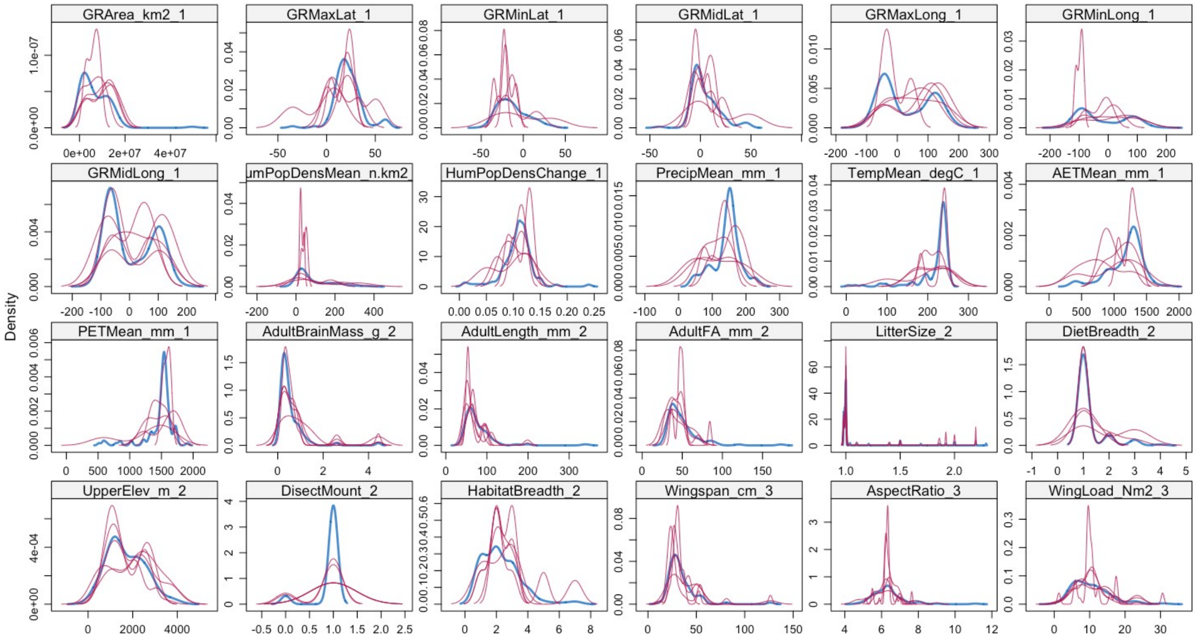


**Figure S2.** Density plots for imputation of missing data for bat traits. Blue lines indicate the distribution of non-missing data, red lines represent distribution of imputed values.

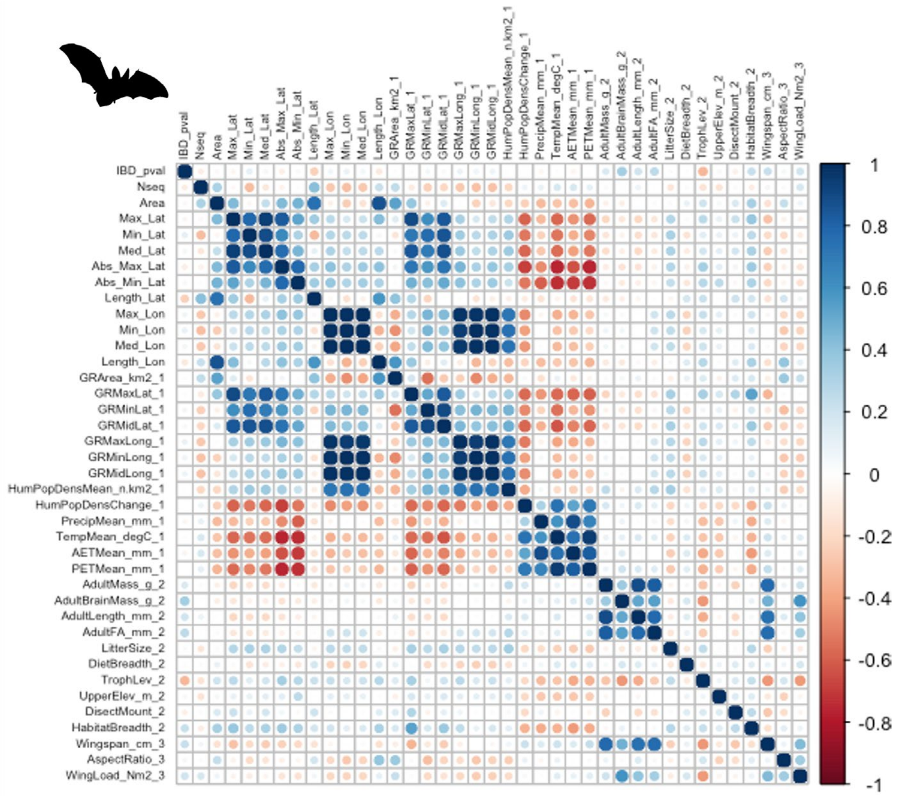


**Figure S3.** Correlation plot of trait variables in the bat dataset.

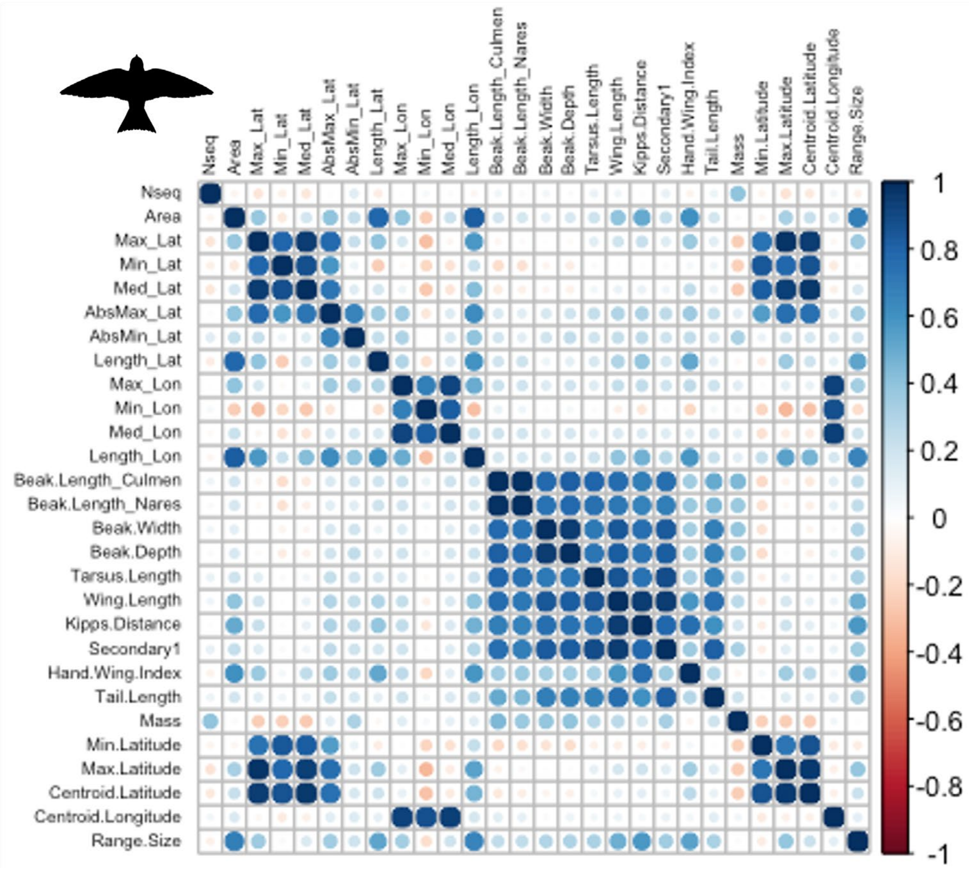


**Figure S4.** Correlation plot of trait variables in the bird dataset.

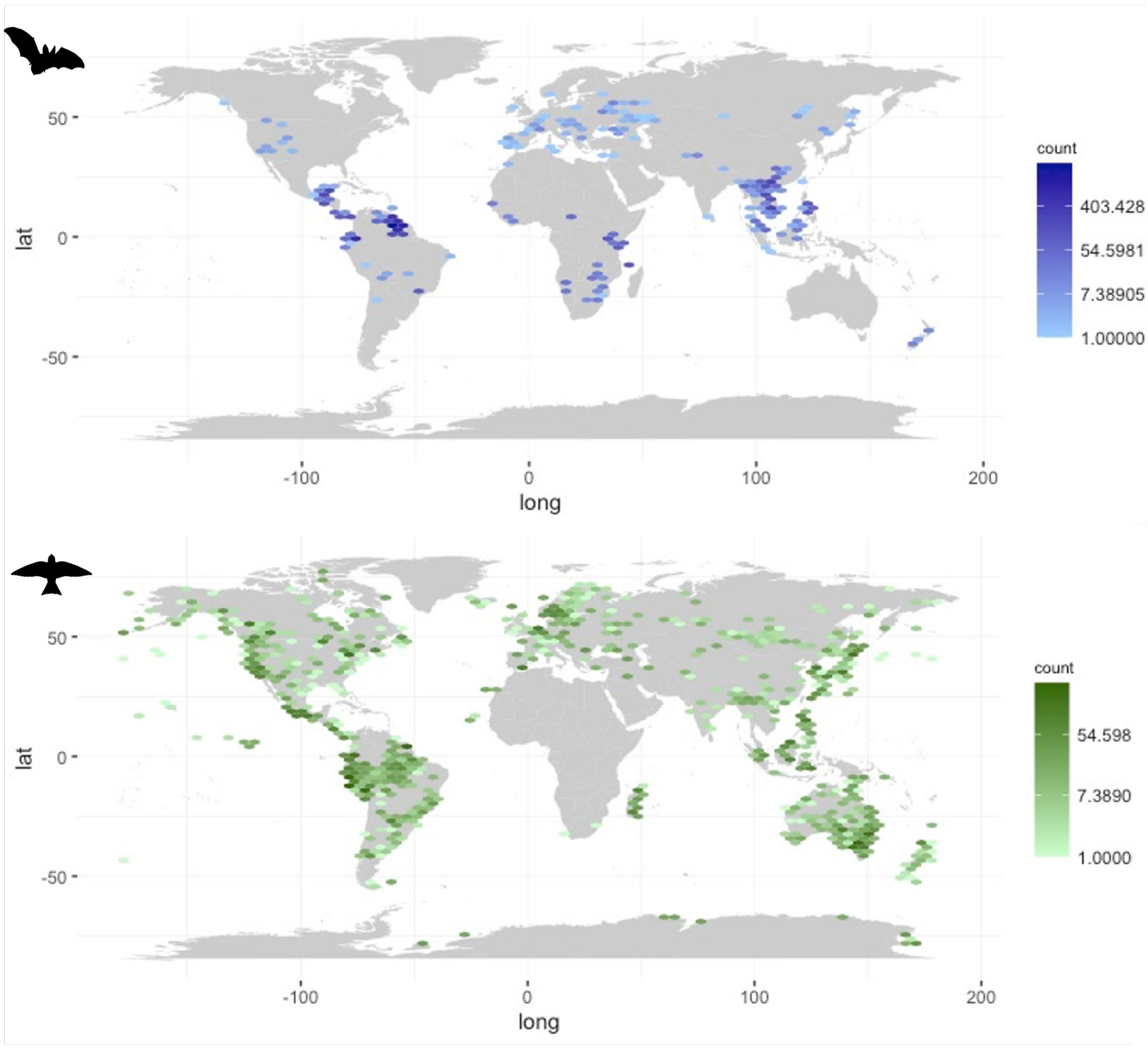


**Figure S5.** Maps of sampling localities and sampling density for bat (top) and bird (bottom) sequences used in the analysis.

**Table S1.** Predictor variables used in the Chiroptera Random Forests classifier with Mean Decrease in Accuracy averaged over 50 independent replicates (AvgMDA), Mean Decrease in Gini averaged over 50 independent replicates (AvgMDG), group means for species estimated to not contain a phylogeographic break (Mean NB), group means for species estimated to contain a break (Mean YB), Welch's two sample t-test statistic (T.Stat), and p-value for t-test (P.Value). Trait variables are listed from highest MDA to lowest, and variables that are significantly different between groups are in bold.

| Label Name | Avg MDA | Avg MDG | MeanB | Mean YB | T.Stat | P.Value (t-test) | MWU Stat | P.Value (MWU) |
|---|---|---|---|---|---|---|---|---|
| **Occurrence Area** | **13.64** | **3.35** | **72.80** | **175.30** | **-3.12** | **0.002** | **1093** | **0.000** |
| Cryptic Diversity Predicted | 12.75 | 1.49 | NA | NA | NA | NA | NA | NA |
| **Latitudinal Range** | **12.61** | **3.33** | **8.30** | **14.20** | **-3.91** | **0.000** | **1108** | **0.000** |
| **Maximum Latitude** | **12.12** | **3.05** | **11.00** | **18.02** | **-2.82** | **0.006** | **1270** | **0.001** |
| Range Bisected by Mountain | 11.10 | 0.84 | NA | NA | NA | NA | NA | NA |
| **Brain Mass** | **9.75** | **2.65** | **0.69** | **0.46** | **2.01** | **0.048** | **2504** | **0.009** |
| Minimum Longitude | 6.09 | 2.29 | 0.66 | -12.37 | 0.85 | 0.397 | 2307.5 | 0.101 |
| **Wingspan** | **5.12** | **2.25** | **39.11** | **34.20** | **2.03** | **0.045** | **2410** | **0.032** |

| | | | | | | | | |
|---|---|---|---|---|---|---|---|---|
| Wing Loading | 4.46 | 2.14 | 10.87 | 9.74 | 1.29 | 0.200 | 2233.5 | 0.201 |
| Trophic Level - Carnivore | 4.36 | 0.64 | NA | NA | NA | NA | NA | NA |
| Human Population Density Change | 4.09 | 1.74 | 0.11 | 0.11 | 0.88 | 0.380 | 2246 | 0.175 |
| Average Evapotranspiration Rate | 3.22 | 2.34 | 11154.4 | 1201.49 | -0.98 | 0.332 | 1871 | 0.623 |
| Mass | 2.79 | 2.12 | 39.52 | 19.04 | 1.55 | 0.126 | 2366.5 | 0.054 |
| Hipposideridae | 2.55 | 0.25 | NA | NA | NA | NA | NA | NA |
| Phyllostomidae | 2.38 | 0.38 | NA | NA | NA | NA | NA | NA |
| Rhinolophidae | 2.17 | 0.28 | NA | NA | NA | NA | NA | NA |
| Realm - Indomalayan/Palearctic | 2.04 | 0.22 | NA | NA | NA | NA | NA | NA |
| Miniopteridae | 2.01 | 0.27 | NA | NA | NA | NA | NA | NA |
| Pteropodidae | 1.83 | 0.37 | NA | NA | NA | NA | NA | NA |
| Trophic Level - Omnivore | 1.62 | 0.37 | NA | NA | NA | NA | NA | NA |
| Realm - Neotropical | 1.42 | 0.40 | NA | NA | NA | NA | NA | NA |
| Realm - Australasian | 0.00 | 0.05 | NA | NA | NA | NA | NA | NA |
| Realm - Nearctic | 0.00 | 0.04 | NA | NA | NA | NA | NA | NA |
| Megadermatidae | 0.00 | 0.05 | NA | NA | NA | NA | NA | NA |
| Mormoopidae | 0.00 | 0.04 | NA | NA | NA | NA | NA | NA |
| Thyropteridae | 0.00 | 0.04 | NA | NA | NA | NA | NA | NA |
| Vespertilionidae | -1.26 | 0.31 | NA | NA | NA | NA | NA | NA |
| Realm - Indomalayan | -1.92 | 0.29 | NA | NA | NA | NA | NA | NA |
| Realm - Australasian/ Indomalayan | -2.19 | 0.17 | NA | NA | NA | NA | NA | NA |
| Realm - Palearctic | -2.59 | 0.21 | NA | NA | NA | NA | NA | NA |
| Realm - Afrotropical/Palearctic | -3.28 | 0.08 | NA | NA | NA | NA | NA | NA |
| Molossidae | -3.31 | 0.16 | NA | NA | NA | NA | NA | NA |
| Realm - Neotropical/Palearctic | -3.90 | 0.13 | NA | NA | NA | NA | NA | NA |
| Noctilionidae | -4.41 | 0.13 | NA | NA | NA | NA | NA | NA |

**Table S2.** Predictor variables used in the Aves Random Forests classifier with Mean Decrease in Accuracy averaged over 50 independent replicates (AvgMDA), Mean Decrease in Gini averaged over 50 independent replicates (AvgMDG), group means for species estimated to not contain a phylogeographic break (Mean NB), group means for species estimated to contain a break (Mean YB), Welch's two sample t-test statistic (T.Stat), and p-value for t-test (P.Value). Trait variables are listed from highest MDA to lowest, and variables that are significantly different between groups are in bold.

| Variable Name | Avg MDA | Avg MDG | Mean NB | Mean YB | T.Stat | P.Value (t-test) | MWU stat | P.Value (MWU) |
|---|---|---|---|---|---|---|---|---|
| **Absolute Maximum Latitude** | **9.71** | **2.68** | **45.84** | **33.27** | **3.82** | **0.000** | **7185** | **0.001** |
| **Beak Length from Nares** | **7.96** | **2.35** | **18.17** | **13.36** | **2.30** | **0.022** | 5789 | 0.7625 |
| **Beak Length from Culmen** | **7.88** | **2.44** | **28.22** | **21.18** | **2.49** | **0.014** | 5776 | 0.7846 |
| Migratory | 7.65 | 0.80 | NA | NA | NA | NA | NA | NA |

| | | | | | | | | |
|---|---|---|---|---|---|---|---|---|
| Procellariiformes | 7.51 | 0.16 | NA | NA | NA | NA | NA | NA |
| Length of Secondary Feather | 7.47 | 2.41 | 85.70 | 72.98 | 1.96 | 0.052 | 6055 | 0.3717 |
| **Wing Length** | **6.76** | **2.37** | **128.31** | **101.75** | **2.22** | **0.028** | 6402 | 0.096 |
| Granivore | 6.67 | 0.46 | NA | NA | NA | NA | NA | NA |
| Invertivore | 6.67 | 0.86 | NA | NA | NA | NA | NA | NA |
| **Mass** | **6.44** | **2.24** | **647.05** | **89.12** | 1.87 | 0.064 | **6578** | **0.040** |
| Insessorial | 6.29 | 0.88 | NA | NA | NA | NA | NA | NA |
| Semi-Open Habitat | 5.96 | 0.75 | NA | NA | NA | NA | NA | NA |
| **Hand-wing Index** | **5.37** | **2.29** | **26.75** | **24.52** | 1.39 | 0.167 | **6598** | **0.036** |
| Forest Habitat | 5.36 | 0.84 | NA | NA | NA | NA | NA | NA |
| Passeriformes | 5.15 | 0.51 | NA | NA | NA | NA | NA | NA |
| Charadriiformes | 5.14 | 0.25 | NA | NA | NA | NA | NA | NA |
| **Tarsus Length** | **5.08** | **2.20** | **30.75** | **23.05** | **3.10** | **0.002** | 6482.5 | 0.065 |
| Centroid Longitude | 5.04 | 2.41 | 5.05 | 18.30 | -1.10 | 0.274 | 5005 | 0.1505 |
| Caprimulgiformes | 4.73 | 0.15 | NA | NA | NA | NA | NA | NA |
| **Beak Depth** | **4.37** | **2.16** | **7.78** | **6.09** | **2.43** | **0.016** | **6665.5** | **0.024** |
| **Kipp's Distance** | **4.23** | **2.20** | **42.32** | **28.65** | **2.22** | **0.027** | 6381.5 | 0.106 |
| Wetland Habitat | 4.13 | 0.38 | NA | NA | NA | NA | NA | NA |
| Beak Width | 4.01 | 2.11 | 6.39 | 5.44 | 1.78 | 0.077 | 6062.5 | 0.363 |
| Pelecaniformes | 3.89 | 0.08 | NA | NA | NA | NA | NA | NA |
| **Absolute Minimum Latitude** | **3.84** | **2.23** | **32.31** | **26.54** | **3.00** | **0.000** | **6824.5** | **0.001** |
| Aquatic | 3.67 | 0.17 | NA | NA | NA | NA | NA | NA |
| Strigiformes | 3.01 | 0.21 | NA | NA | NA | NA | NA | NA |
| Generalist | 2.99 | 0.47 | NA | NA | NA | NA | NA | NA |
| Anseriformes | 2.56 | 0.04 | NA | NA | NA | NA | NA | NA |
| Terrestrial | 2.16 | 0.52 | NA | NA | NA | NA | NA | NA |
| Marine Habitat | 1.89 | 0.17 | NA | NA | NA | NA | NA | NA |
| Partially Migratory | 1.78 | 0.41 | NA | NA | NA | NA | NA | NA |
| Human Modified Habitat | 1.60 | 0.26 | NA | NA | NA | NA | NA | NA |
| Trophic Level - Herbivore | 1.54 | 0.38 | NA | NA | NA | NA | NA | NA |
| Tail Length | 0.67 | 2.12 | 82.64 | 72.49 | 1.54 | 0.124 | 6351.5 | 0.121 |
| Open Habitat | 0.58 | 0.38 | NA | NA | NA | NA | NA | NA |
| Nectarivore | 0.20 | 0.13 | NA | NA | NA | NA | NA | NA |
| Trophic Level - Omnivore | 0.00 | 0.40 | NA | NA | NA | NA | NA | NA |
| Cuculiformes | 0.00 | 0.07 | NA | NA | NA | NA | NA | NA |
| Falconiformes | 0.00 | 0.03 | NA | NA | NA | NA | NA | NA |
| Gruiformes | 0.00 | 0.03 | NA | NA | NA | NA | NA | NA |
| Psittaciformes | 0.00 | 0.03 | NA | NA | NA | NA | NA | NA |
| Suliformes | 0.00 | 0.02 | NA | NA | NA | NA | NA | NA |
| Omnivore | 0.00 | 0.39 | NA | NA | NA | NA | NA | NA |
| Range Size | -0.06 | 2.10 | $1.2 \times 10^7$ | $1.1 \times 10^7$ | 0.33 | 0.744 | NA | NA |

| | | | | | | | | |
|---|---|---|---|---|---|---|---|---|
| Vertivore | -0.10 | 0.15 | NA | NA | NA | NA | NA | NA |
| Sampling Area | -0.15 | 2.13 | 1419.95 | 1493.73 | -0.19 | 0.847 | 5755.5 | 0.820 |
| Shrubland Habitat | -0.29 | 0.36 | NA | NA | NA | NA | NA | NA |
| Riverine Habitat | -1.31 | 0.12 | NA | NA | NA | NA | NA | NA |
| Sphenisciformes | -1.32 | 0.07 | NA | NA | NA | NA | NA | NA |
| Coraciiformes | -1.52 | 0.14 | NA | NA | NA | NA | NA | NA |
| Woodland Habitat | -1.82 | 0.30 | NA | NA | NA | NA | NA | NA |
| Piciformes | -1.93 | 0.16 | NA | NA | NA | NA | NA | NA |
| Frugivore | -1.98 | 0.16 | NA | NA | NA | NA | NA | NA |
| Herbivore | -2.05 | 0.08 | NA | NA | NA | NA | NA | NA |
| Grassland Habitat | -2.19 | 0.22 | NA | NA | NA | NA | NA | NA |
| Galliformes | -3.73 | 0.12 | NA | NA | NA | NA | NA | NA |

**Table S3.** Random Forests model results for classifiers built using significant isolation by distance as the response variable.

| | OOB | Class (N) | Class (Y) | Sensitivity | Specificity | Precision |
|---|---|---|---|---|---|---|
| Bats | 0.2862 | 0.7388 | 0.0956 | 0.6831 | 0.8462 | 0.3333 |
| Birds | 0.4065 | 0.6557 | 0.2372 | 0.8947 | 0.250 | 0.6538 |

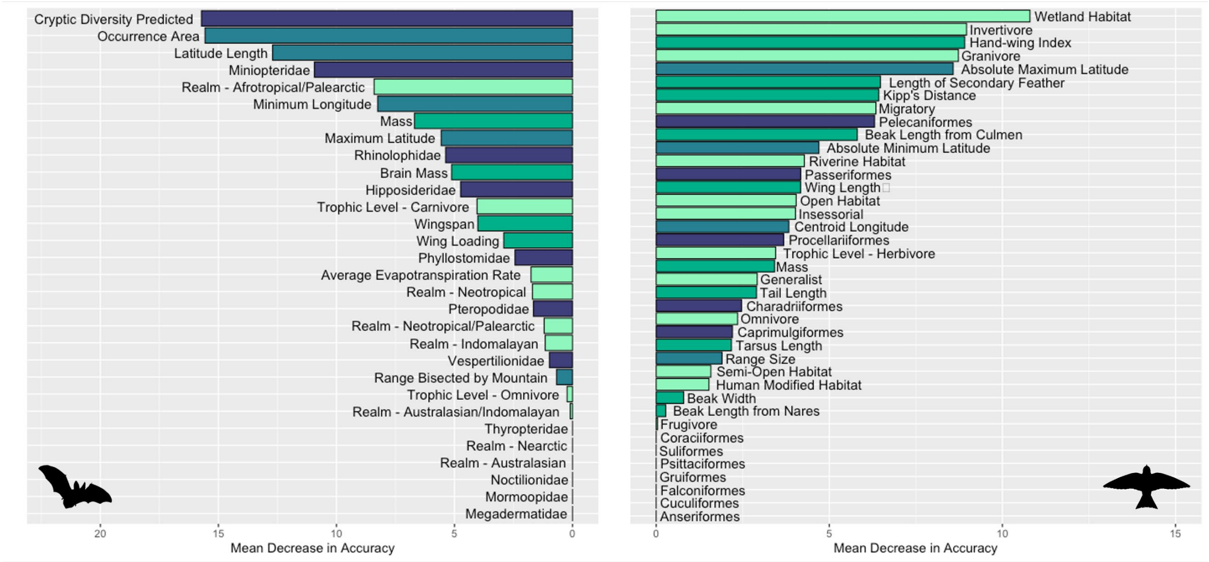


**Figure S6.** Variable importance plot for random forests classifier using isolation by distance result as the response variable for bats (left) and birds (right). Variable importance is measured by mean decrease in accuracy.
